# Single mutation at a highly conserved region of chloramphenicol acetyltransferase enables thermophilic isobutyl acetate production directly from cellulose by *Clostridium thermocellum*

**DOI:** 10.1101/756916

**Authors:** Hyeongmin Seo, Jong-Won Lee, Sergio Garcia, Cong T. Trinh

## Abstract

**Background:** Esters are versatile chemicals and potential drop-in biofuels. To develop a sustainable production platform, microbial ester biosynthesis using alcohol acetyltransferases (AATs) has been studied for decades. Volatility of esters endows thermophilic production with advantageous downstream product separation. However, due to the limited thermal stability of AATs known, the ester biosynthesis has largely relied on use of mesophilic microbes. Therefore, developing thermostable AATs is important for thermophilic ester production directly from lignocellulosic biomass by the thermophilic consolidated bioprocessing (CBP) microbes, e.g., *Clostridium thermocellum*.

**Results:** In this study, we engineered a thermostable chloramphenicol acetyltransferase from *Staphylococcus aureus* (CAT_Sa_) for enhanced isobutyl acetate production at elevated temperature. We first analyzed the broad alcohol substrate range of CAT_Sa_. Then, we targeted a highly conserved region in the binding pocket of CAT_Sa_ for mutagenesis. The mutagenesis revealed that F97W significantly increased conversion of isobutanol to isobutyl acetate. Using CAT_Sa_ F97W, we demonstrated the engineered *C. thermocellum* could produce isobutyl acetate directly from cellulose.

**Conclusions:** This study highlights that CAT is a potential thermostable AAT that can be harnessed to develop the thermophilic CBP microbial platform for biosynthesis of designer bioesters directly from lignocellulosic biomass.

## Introduction

Esters are versatile chemicals which have been used as lubricants, solvents, food additives, fragrances and potential drop-in fuels [1]. Currently, ester production largely relies on synthesis from petroleum or extraction from plants, which makes it neither sustainable nor economically feasible. Therefore, microbial production of esters has been studied for decades [2–7]. Most studies have employed an alcohol acetyltransferase (E.C. 2.3.1.84, AAT), belonging to a broad acetyltransferase class, that can synthesize a carboxylic ester by condensing an alcohol and an acyl-CoA in a thermodynamically favorable aqueous environment [5]. For example, an *Escherichia coli,* engineered to use this biosynthetic pathway, could achieve high titer of isobutyl acetate [6, 7]. With appropriate expression of AATs and availability of alcohol and acyl-CoA moieties, various types of esters can be produced [2, 4]. Due to high volatility of esters, ester production at elevated temperature can benefit downstream product separation and hence reduce the process cost. Interestingly, it has recently been shown that for the same total carbon chain length, short-chain esters are less toxic to microbial health than alcohols, which is potentially beneficial for ester fermentation [8]. However, most of the AATs known to date are isolated from mesophilic microbes or plants, and none of them has been reported to be active at elevated temperatures (> 50°C). The highest temperature reported for ester production is 42°C in a thermotolerant yeast [9]. Hence, finding and developing a thermostable AAT is crucial to produce esters at elevated temperature.

Chloramphenicol acetyltransferase (E.C. 2.3.1.28, CAT) is another acetyltransferase class that has been found in various microbes [10]. This enzyme acetylates chloramphenicol, a protein synthesis inhibitor, by transferring the acetyl group from acetyl-CoA. The acetylation of chloramphenicol detoxifies the antibiotic compound and confers chloramphenicol resistance in bacteria. Recent studies have implied that CATs likely recognize a broad substrate range for alcohols and acyl-CoAs [7]. In addition, high thermal stability of some CATs enables them to be used as selection markers in thermophiles [11–13]. Therefore, CAT can function or be repurposed as a thermostable AAT suitable for ester biosynthesis at elevated temperature.

In this study, we engineered a CAT from *Staphylococcus aureus* (CAT_Sa_) for thermophilic isobutyl acetate production. First, we investigated a broad alcohol substrate range of CAT_Sa_. Protein homology modeling along with sequence alignment were performed to identify the binding pocket of CAT_Sa_ as a potential target for protein engineering to enhance condensation of isobutanol and acetyl-CoA. *In silico* mutagenesis successfully discovered a variant (F97W) of CAT_Sa_ that was then experimentally validated for improved catalytic activity towards isobutanol. As a proof of concept, the engineered CAT_Sa_ was successfully expressed in *Clostridium thermocellum.* We demonstrated the F97W CAT_Sa_-overexpressing for consolidated bioprocessing (CBP) to produce isobutyl acetate directly from cellulose without a need for external supply of cellulases. To our knowledge, this study presents the first demonstration of CAT engineering to enable thermophilic ester production directly from cellulose.

## Results and discussion

### *In silico* and rapid *in vivo* characterization of a thermostable chloramphenicol acetyltransferase(s) for broad alcohol substrate range

To develop a thermophilic microbial ester production platform, a thermostable AAT is required. Unfortunately, the AATs known to date are isolated from mesophilic yeasts or plants, and none of them has been reported to be active at a temperature above 50°C. To tackle this problem, we chose CATs to investigate their potential functions as a thermostable AAT because some thermophilic CATs have been successfully used as a selection marker in thermophiles [13–17] and others have been shown to perform the acetylation for not only chloramphenicol but various alcohols like AATs [18–21] [7] (Figure 1A, S1A). As a proof-of-study, we investigated CAT_Sa_, classified as Type A-9, from the plasmid pNW33N for a broad range of alcohol substrates as it has been widely used for genetic engineering in *C. thermocellum* at elevated temperature (≥ 50°C) [13–15].

**Figure 1.**
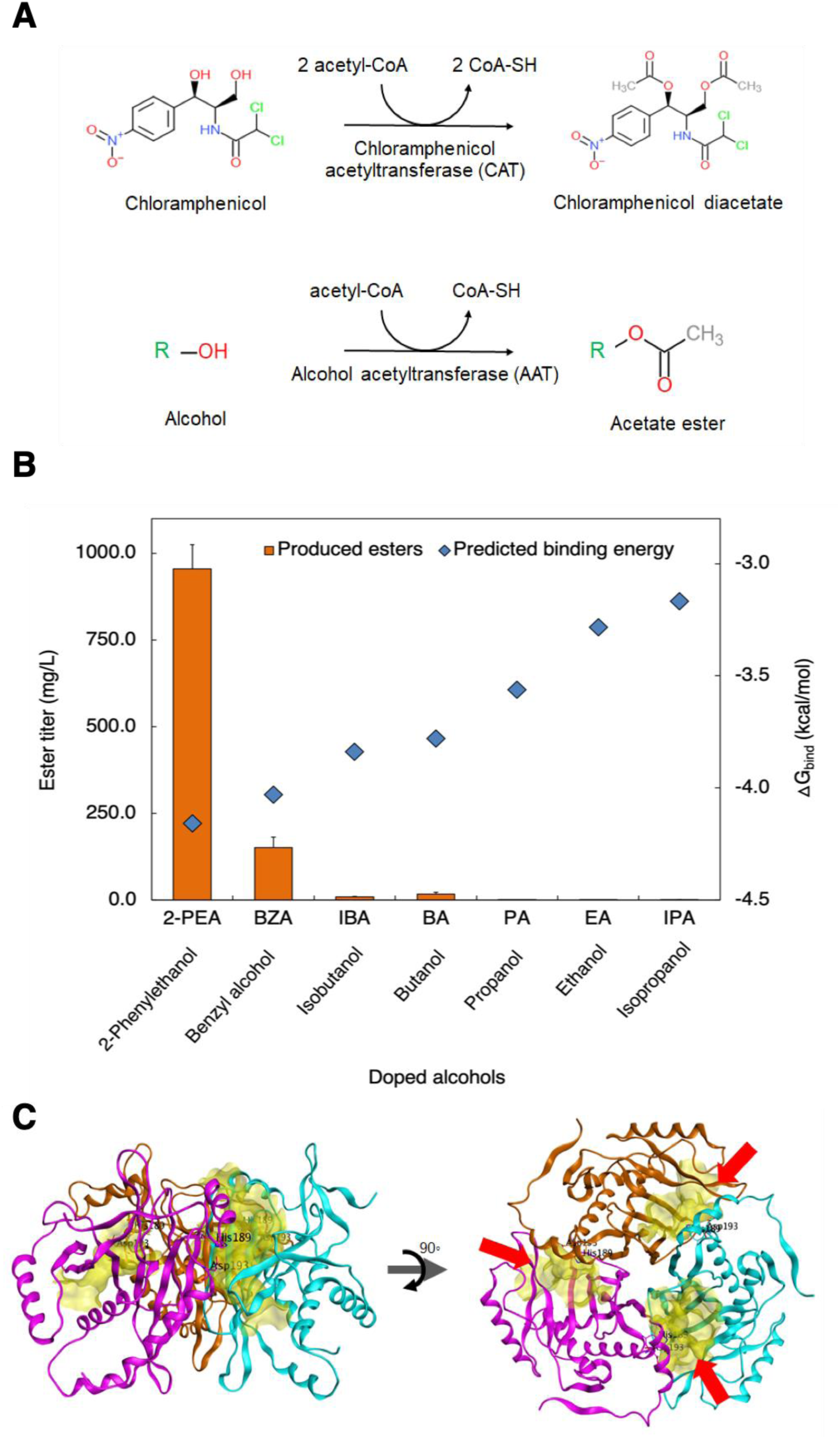
Broad substrate specificity of CAT_Sa_. **(A)** Acetylation of chloramphenicol and alcohol by a chloramphenicol acetyltransferase (CAT) and an alcohol acetyltransferase (AAT), respectively. **(B)** Comparison between the predicted binding free energies for various alcohols bound to the binding pocket of CAT_Sa_ and the titer of esters produced by the CAT_Sa_-overexpressing *E. coli* with external supply of alcohols. **(C)** Structure of the CAT_Sa_ homology model. The red arrows indicate the binding pockets formulated by the trimeric structure of CAT_Sa_.

We first conducted alcohol docking simulations using the homology model. Remarkably, the model predicted binding affinities of short-to-medium chain length alcohols (e.g., ethanol, propanol, isopropanol, butanol, and isobutanol) and aromatic alcohols (e.g., benzyl alcohol and phenethyl alcohol) to the binding pocket. The change in the protein’s Gibbs free energy upon the substrate binding was ordered as follows: 2-phenethyl alcohol > benzyl alcohol > isobutanol > butanol > propanol > ethanol > isopropanol (Figure 1B).

To quickly evaluate the *in silico* docking simulation results experimentally, we next performed *in vivo* characterization of a CAT_Sa_-overexpressing *E. coli* and screened for acetate esters production. Acetyl-CoA was derived from glycolysis while various alcohols were externally supplied to the media. Remarkably, the results exhibited the same trends of specificities of CAT_Sa_ towards alcohols as predicted by the *in silico* docking simulation (Figure 1B). The CAT_Sa_-overexpressing *E. coli* produced all the expected acetate esters including ethyl acetate, propyl acetate, isopropyl acetate, butyl acetate, isobutyl acetate, benzyl acetate, and 2-phenethyl acetate at titers of 1.12 ± 0.07, 2.30 ± 0.28, 0.08 ± 0.02, 9.75 ± 1.57, 17.06 ± 6.04, 152.44 ± 29.50, and 955.27 ± 69.50 mg/L and specific ester production rates of 0.02 ± 0.00, 0.05 ± 0.01, 0.00 ± 0.00, 0.19 ± 0.03, 0.34 ± 0.12, 3.02 ± 0.57, and 19.27 ± 1.32 mg/gDCW/h, respectively. We observed that the specific ester production titers and rates are higher for aromatic alcohols than linear, short-chain alcohols likely because the hydrophobic binding pocket of CAT_Sa_ has been evolved towards chloramphenicol [22], an aromatic antibiotic (Figure 1C). Specifically, the bulky binding pocket of CAT_Sa_ likely contributes to more interaction with the aromatic substrates than the short, linear-chain alcohols (Figure S1B and S1C).

Overall, thermostable CATs, e.g., CAT_Sa_, can have broad range of substrate specificities towards linear, short-chain, and aromatic alcohols and hence can be harnessed as AATs for novel ester biosynthesis at elevated temperature.

### Discovery of a CAT_Sa_ variant improving conversion isobutanol and acetyl CoA into isobutyl acetate

Since the *in vivo* activity of CAT_Sa_ is more than 50-fold higher for the aromatic alcohols than isobutanol, we asked whether its activity could be improved for isobutyl acetate biosynthesis. Using the *in silico* analysis, we started by examining whether any modification of the binding pocket of CAT_Sa_ could improve the activity towards isobutanol. According to the homology model, the binding pocket consists of Tyr-20, Phe-27, Tyr-50, Thr-88, Ile-89, Phe-90, Phe-97, Ser-140, Leu-141, Ser-142, Ile-143, Ile-144, Pro-145, Trp-146, Phe-152, Leu-154, Ile-166, Ile-167, Thr-168, His-189, Asp-193, Gly-194, and Tyr-195, where the His189 and Asp193 are the catalytic sites (Figure 2A). Since chloramphenicol resistance is likely a strong selective pressure throughout evolution, we expected all CATs to exhibit a common binding pocket structure. Unsurprisingly, conserved sequences in the binding pocket were observed by protein sequence alignment of CAT_Sa_ with other CATs of Type A (Figure S2A). Especially, Pro-85 and Phe-97 were highly conserved in CATs of not only Type-A but also Type-B (Figure 2B and Figure S2B).

**Figure 2.**
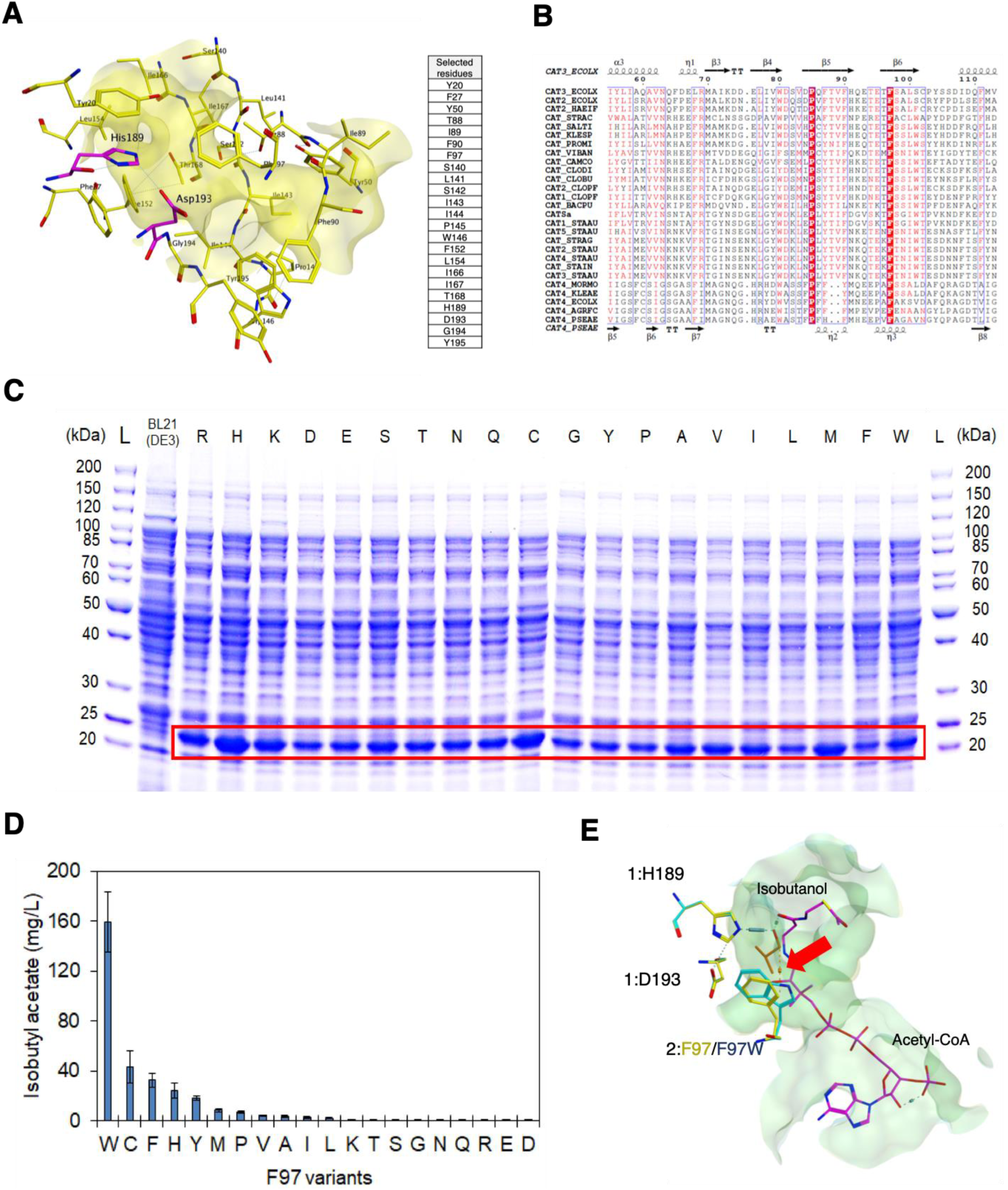
Discovery of CAT_Sa_ F97W responsible for enhanced activity towards isobutanol. **(A)** A binding pocket of CAT_Sa_ and associated amino acid residues. The catalytic site residues are in purple. **(B)** Protein sequence alignment of CAT_Sa_ with different CATs. **(C)** SDS-PAGE analysis of soluble fractions of CAT_Sa_ F97 variants. The soluble fractions of overexpressed CAT_Sa_ F97 variants are shown in the red box. **(D)** Screening of F97 site-saturated mutagenized variants for enhanced isobutyl acetate production in *E. coli*. The letters indicate amino acids substituting F in the wildtype CAT_Sa_. **(E)** Superposed binding pocket structure of the wildtype and CAT_Sa_ F97W mutant. The red arrow indicates a CH-π interaction between the hydrogen of isobutanol and the indole ring of F97W.

Based on the binding pocket identified, we performed docking simulation with alanine and residue scans using the acetyl-CoA-isobutanol-CAT_Sa_ complex to identify potential candidates for mutagenesis (Figure S3A and S3B). Remarkably, the top three variant candidates were suggested at the Phe-97 residue. This residue is involved in the formation of a tunnel-like binding pocket [22]. Motivated by the analysis, Phe-97 was chosen for site saturated mutagenesis, and the variants were screened in *E. coli* for isobutyl acetate production by external supply of isobutanol.

The result showed that all the F97 variants did not affect protein expression levels in *E. coli* (Figure 2C). Among the variants characterized, the F97W variant exhibited the best performance (Figure 2D). As compared to the wildtype, the F97W variant enhanced the isobutyl acetate production by 4-fold. Subsequent *in silico* analysis showed that the mutation created a CH-π interaction between the hydrogen of isobutanol and the indole ring of F97W (Figure 2E). The model also indicated no change in distance between the isobutanol and active site (His-189) in F97W. Therefore, the CH-π interaction is likely responsible for the improved activity of F97W variant towards isobutyl acetate biosynthesis.

### *In vitro* Characterization of CAT_Sa_ F97W

Before deploying CAT_Sa_ F97W for isobutyl acetate biosynthesis in the thermophile CBP organism *C. thermocellum*, we checked whether the F97W mutation affected thermal stability of the enzyme. We overexpressed and purified both the wildtype CAT_Sa_ and CAT_Sa_ F97W variant (Figure 3A). The SDS-PAGE analysis confirmed the expression and purification of the enzymes by bands with the expected monomer size (25.8 kDa). Thermofluor assay revealed that the F97W variant slightly lowered the wildtype melting point from 72°C to 68.3°C (Figure 3B). Since CAT_Sa_ F97W maintained high melting point, it is possible that CAT_Sa_ F97W still maintains its functionality at high temperature (≥ 50°C) but needs to be thoroughly characterized.

**Figure 3.**
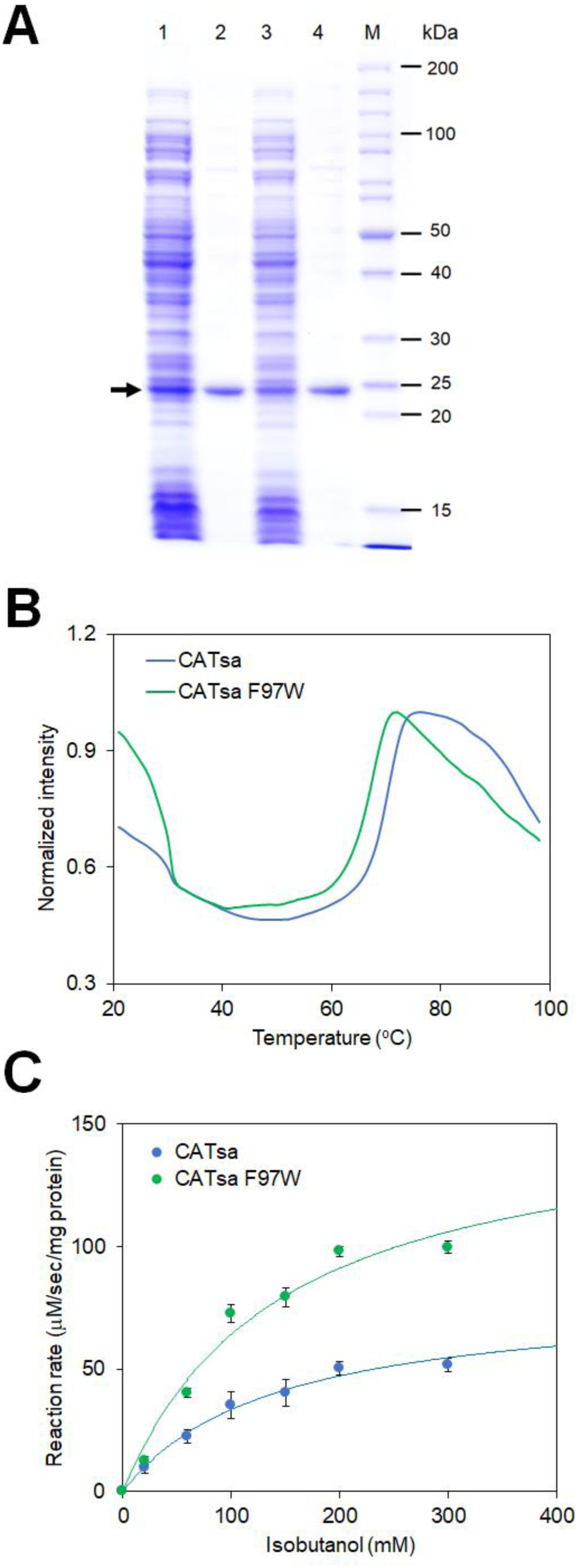
*In vitro* characterization of the wildtype CAT_Sa_ and CAT_Sa_ F97W mutant. **(A)** SDS-PAGE of the purified CAT_Sa_ and CAT_Sa_ F97W. The black arrow indicates the expected size of expressed target proteins, including CAT_Sa_ and CAT_Sa_ F97W. Notations: column 1, crude cell extract of IPTG induced *E. coli* BL21(DE3) harboring pET_CAT_Sa_; column 2, His-tag purified CAT_Sa_; column 3, crude extract of IPTG induced *E. coli* BL21(DE3) harboring pET_ CAT_Sa_ F97W; column 4, His-tag purified CAT_Sa_ F97W; and M, protein ladder. **(B)** Melting curve of CAT_Sa_ and CAT_Sa_ F97W. The intensity was normalized by each maximum value. **(C)** Michaelis-Menten plots of CAT_Sa_ and CAT_Sa_ F97W for various isobutanol concentrations at 50°C. The co-substrate, acetyl-CoA, was supplemented at the saturated concentration of 2 mM. The error bars represent standard deviation of three biological replicates.

Table 2 shows the *in vitro* enzymatic activities of both the wildtype CAT_Sa_ and CAT_Sa_ F97W at 50°C. The turnover number (kcat) of CAT_Sa_ F97W was two times higher than that of the wildtype. The increased turnover number of CAT_Sa_ F97W led to 1.9-fold increase in enzymatic efficiency (kcat/K_M_, 4.08 ± 0.62, 1/M/sec) while the mutation did not result in significant change in K_M_. The improved enzymatic efficiency of CAT_Sa_ F97W agrees with the enhanced isobutanol production observed in the *in vivo* characterization using the CAT_Sa_-overexpressing *E. coli* (Figure 2C).

Based on the rigidity of the binding pocket, we originally presumed that mutagenesis on the binding pocket would result in activity loss towards chloramphenicol. Surprisingly, CAT_Sa_ F97W retained the activity towards chloramphenicol (Table 2). The F97W mutation decreased kcat but also lowered K_M_, resulting in a compensation effect. Turnover number of CAT_Sa_ (kcat, 202.97 ± 3.36, 1/sec) was similar to the previously reported value by Kobayashi *et al.* [12], but K_M_ (0.28 ± 0.02, mM) was about 1.75-fold higher. The difference might attribute to the experimental condition and analysis performed. Kobayashi *et al.* used chloramphenicol in a range of 0.05-0.2 mM for the assay and the Lineweaver-Burk method for analysis, while we used a 0-1.0 mM range with a nonlinear regression analysis method. Interestingly, affinity towards acetyl-CoA was independent of the alcohol co-substrates (Table S2), suggesting that the alcohol affinity would be likely the main bottleneck for microbial production of isobutyl acetate.

Taken altogether, the F97W mutation not only resulted in 1.9-fold higher enzymatic efficiency towards isobutanol but also retained thermal stability of CAT_Sa_. Thus, CAT_Sa_ F97W variant can serve a starting candidate to demonstrate direct biosynthesis of isobutyl acetate at elevated temperature by *C. thermocellum*.

### Isobutyl acetate production from cellulose at elevated temperature by an engineered *C. thermocellum* overexpressing CAT_Sa_ F97W

We next investigated whether *C. thermocellum* overexpressing CAT_Sa_ F97W could produce isobutyl acetate at elevated temperature. This thermophile was chosen because it has a high cellulolytic activity suitable for CBP, a one-step process configuration for cellulase production, cellulose hydrolysis, and fermentation for direct conversion of lignocellulosic biomass to fuels and chemicals [23]. Furthermore, studies have demonstrated that the wildtype *C. thermocellum* has native metabolism capable of endogenously producing precursor metabolites for ester biosynthesis, such as acetyl-CoA, isobutyryl-CoA, as well as ethanol [24] and higher alcohols (e.g., isobutanol) under high cellulose loading fermentation [25–27] (Figure 4A, S5A).

**Figure 4.**
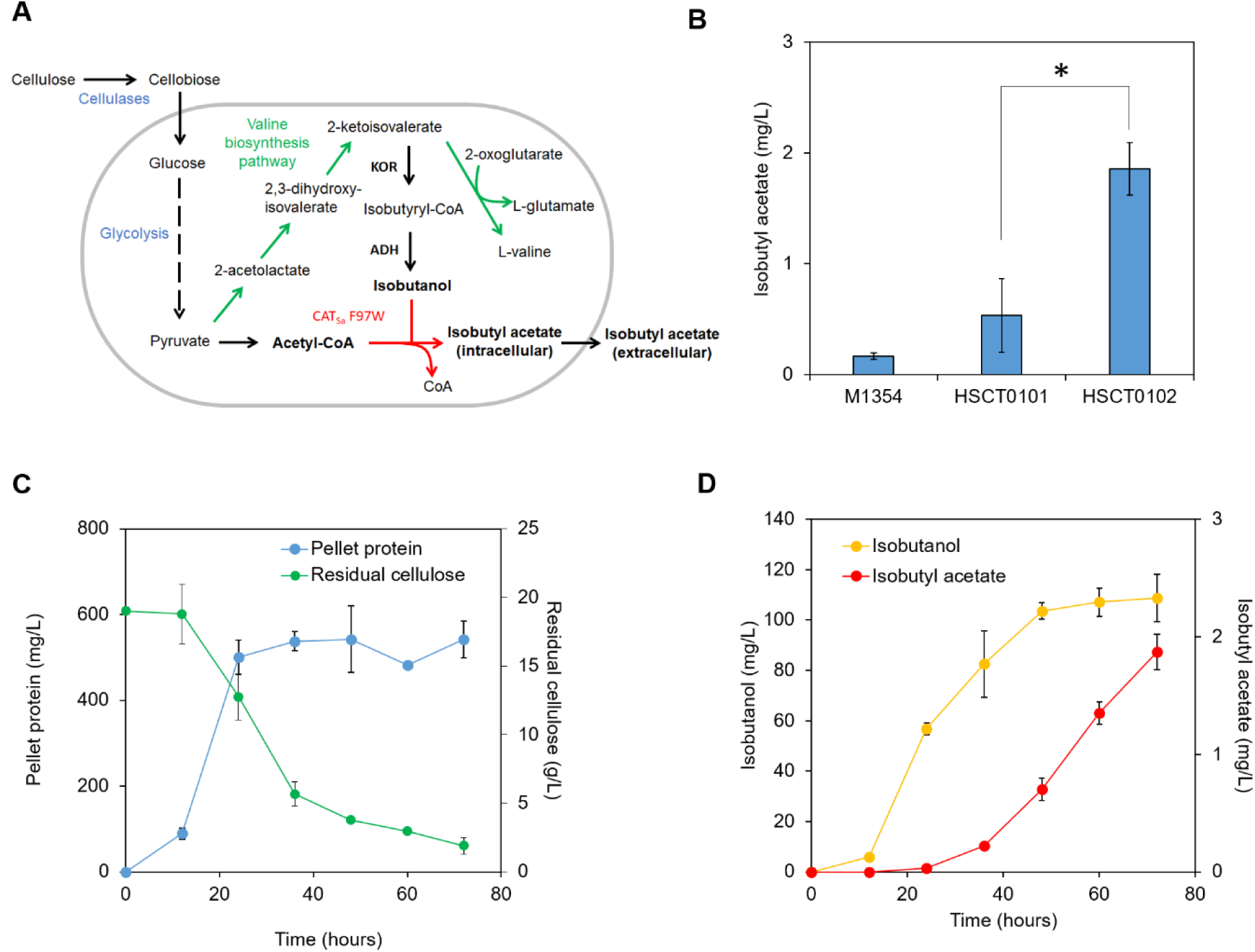
Isobutyl acetate production in the engineered *C. thermocellum*. **(A)** A simplified isobutyl acetate production pathway from cellulose in *C. thermocellum*. **(B)** Biosynthesis of isobutyl acetate of the wildtype and engineered *C. thermocellum* strains at 55°C from cellobiose with external supply of 2 g/L of isobutanol. Isobutyl acetate was measured after 24 hours from the hexadecane layer of cell cultures. Initial OD of each cell culture was in a range of 0.8−1.0. The error bars represent standard deviation of five biological replicates. Statistical analysis: t-test, “*” p value < 4 × 10^−4^, t = −6.475, df = 7. **(C)** Kinetic profiles of cell growth and residual cellulose of HSCT0102. The error bars represent standard deviation of three biological replicates. **(D)** Kinetic profiles of isobutanol and isobutyl acetate production. The error bars represent standard deviation of three biological replicates. Abbreviation: KOR, 2-ketoisovalerate ferredoxin oxidoreductase; ADH, alcohol dehydrogenase.

We started by generating two isobutyl acetate-producing strains, HSCT0101 and HSCT0102, by introducing the plasmids pHS0024 (harboring the wildtype CAT_Sa_) and pHS0024_F97W (harboring the mutant CAT_Sa_ F97W) into *C. thermocellum* DSM1313. Colonies were isolated on antibiotics selective plates at 55°C. Successful transformation clearly indicated that CAT_Sa_ F97W conferred the thiamphenicol resistance and hence maintained CAT activity. This result agrees with the *in vitro* enzymatic activity of CAT_Sa_ F97W (Table 2).

We next evaluated whether the *C. thermocellum* strains could synthesize isobutyl acetate from cellobiose. Since the endogenous isobutanol production from a typical cellobiose concentration (5 g/L) is low [27], we supplemented the media with 2 g/L isobutanol. Both HSCT0101 and HSCT0102 could produce isobutyl acetate at 55°C as expected. Like the *in vivo* characterization in *E. coli* (Figure 2C), HSCT0102 outperformed HSCT0101 with 3.5-fold increase in isobutyl acetate production (Figure 4B). Interestingly, we also observed the parent *C. thermocellum* M1354 produced a trace amount of isobutyl acetate (< 0.1 mg/L) even though this strain does not harbor a CAT (Figure S4). This phenomenon was only observed when hexadecane overlay was used during fermentation for ester extraction. One possible explanation is the endogenous activity of esterases in *C. thermocellum* might have been responsible for low isobutyl acetate production while the organic phase overlay helps extract the target ester. It should be noted that the esterase reaction is reversible and more thermodynamically favorable for ester degradation than biosynthesis.

Finally, we tested whether HSCT0102 could produce isobutyl acetate directly from cellulose at elevated temperature (55°C) without external supply of isobutanol. After 72 hours, cell mass, containing 550 mg/L of pellet protein, reached 1.04 g/L, and 17 g/L of cellulose were consumed (Figure 4C). About 103 mg/L of isobutanol were produced for the first 48 hours, and further increased up to 110 mg/L for additional 24 hours (Figure 4D). Besides isobutanol, *C. thermocellum* also produced other fermentative metabolites, including ethanol, formate, acetate, and lactate, as expected (Figure S5A, S5B). For the target isobutyl acetate production, HSCT0102 did not produce isobutyl acetate for the first 24 hours but started accumulating the target product for the next 48 hours. The observed profile of isobutyl acetate production could be attributed to the low substrate affinity of CAT_Sa_ F97W (Table 2). The final titer of isobutyl acetate reached 1.9 mg/L, achieving about 0.12% (w/w) cellulose conversion.

Besides the production of the desirable ester isobutyl acetate, we also observed that HSCT0102 produced other detectable esters such as ethyl acetate, ethyl isobutyrate, and isobutyl isobutyrate (Figure S5A, S5C, S5D). Endogenous biosynthesis of these esters could be explained from the complex redox and fermentative metabolism of *C. thermocellum* [26, 28]. *C. thermocellum* can endogenously synthesize the precursor metabolites, acetyl-CoA and ethanol via the ethanol biosynthesis pathway while *C. thermocellum* can endogenously produce the precursor metabolites, isobutyryl-CoA and isobutanol via the valine biosynthesis (Figure S5A). With the availability of four precursor metabolites, *C. thermocellum* could produce ethyl acetate, ethyl isobutyrate, isobutyl acetate, and isobutyl isobutyrate as observed experimentally (Figure S5A, S5C, S5D).

Taken altogether, *C. thermocellum* overexpressing CAT_Sa_ successfully produced the target isobutyl acetate from cellulose at elevated temperature (55°C). The engineered CAT_Sa_ F97W enhanced isobutyl acetate production and is capable of producing other types of esters.

## Conclusions

This study demonstrated that a CAT can function and/or be re-purposed as an AAT for novel biosynthesis of designer esters at elevated temperature. Both *in silico* and *in vivo* characterization discovered a broad alcohol substrate range of the thermostable chloramphenicol acetyltransferase from *Staphylococcus aureus* (CAT_Sa_). Discovery of the F97W mutation of CAT_Sa_ by model-guided protein engineering enhanced isobutyl acetate production. This study presented the first report on the consolidated bioprocessing of cellulose into ester(s) by the thermophilic CBP organism *C. thermocellum* harboring an engineered thermostable CAT_Sa_ F97W. Overall, this research helps establish a foundation for engineering non-model organisms for direct conversion of lignocellulosic biomass into designer bioesters.

## Materials and methods

### Bacterial strains and plasmids

Bacterial strains and plasmids used in this study are listed in Table 1. *Clostridium thermocellum* DSM1313 Δ*hpt* (M1354) strain was used as a host for the thermophilic ester production. It should be noted that the deletion of hypoxanthine phosphoribosyltransferase gene (*hpt*, Clo1313_2927) in the wildtype DSM1313 allows genetic engineering by 8-Azahypoxanthine (8-AZH) counter selection; this deletion does not have any known adverse effect on cell growth and metabolism [29, 30]. The plasmid pNW33N, containing CAT_Sa_, is thermostable and was used to express various CATs in *C. thermocellum*. The pET plasmids were used for molecular cloning and enzyme expression in *E. coli*.

**Table 1.** Plasmids and strains used in this study. The plasmids containing mutagenized genes are presented in Table S1.

| Name | Descriptions | Source |
| --- | --- | --- |
| *Plasmids* |  |  |
| pNW33N | *Bacillus-E. coli* shuttle vector, Cm<sup>R</sup>, pBC1 ori for gram positive strains, pBR322 ori for *E. coli*, source of CAT<sub>Sa</sub> | Bacillus Genetic Stock Center |
| pETDuet-1 | pBR322 ori, Amp<sup>R</sup>, lacI, T7lac promoter | Novagen |
| pET_CAT<sub>Sa</sub> | CAT<sub>Sa</sub> wild type encoding gene between BamHI, SacI site, pETDuet-1 backbone, 6X His-tag at N-terminus | This study |
| pET_CAT<sub>Sa</sub> F97W | F97W site directed variant, pET_CAT<sub>Sa</sub> backbone | This study |
| pHS0024 | CAT<sub>Sa</sub> wild type gene under *C. thermocellum* PgapDH promoter, downstream of Clo1313_2927 for the transcription terminator, tdk operon under cbp promoter substituting with the native cat selection marker, pNW33N plasmid backbone | This study |
| pHS0024_F97W | CAT<sub>Sa</sub> F97W site-directed mutated from pHS0024 | This study |
| *Strains* |  |  |
| *E. coli* Top10 | Host for molecular cloning, *mcrA*, $\Delta(mrr-hsdRMS-mcrBC)$, *Phi80lacZ(del)M15*, $\Delta lacX74$, *deoR*, *recA1*, *araD139*, $\Delta(ara-leu)7697$, *galU*, *galK*, *rpsL(SmR)*, *endA1*, *nupG* | Invitrogen |
| *E. coli* BL21 (DE3) | *E. coli* B *dcm*, *ompT*, *hsdS(rB-mB-)*, *gal* | Invitrogen |
| M1354 | *C. thermocellum* DSM1313 $\Delta hpt$ | [29] |
| HSCT0101 | M1354 harboring pHS0024 | This study |
| HSCT0102 | M1354 harboring pHS0024_F97W | This study |

**Table 2.**
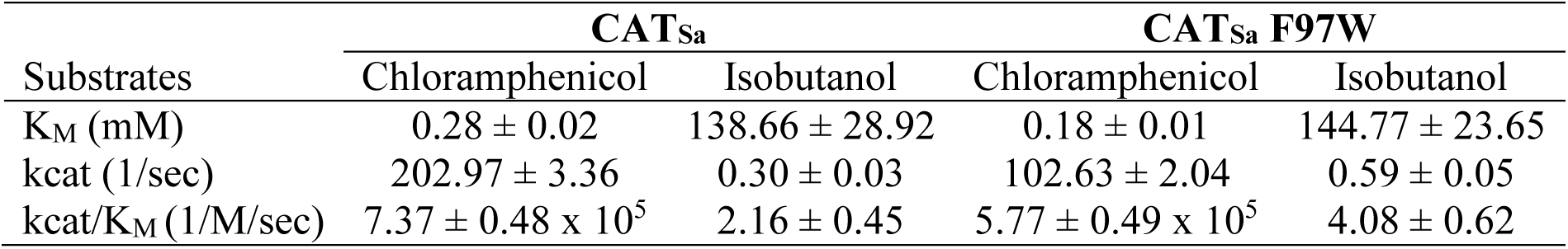
Kinetic parameters of the wildtype CAT_Sa_ and mutant CAT_Sa_ F97W. The reactions were performed at 50°C. The co-substrate, acetyl-CoA, was supplied at the saturated concentration of 2 mM. Tm of CAT_Sa_ and CAT_Sa_ F97W are 72.0 ± 0.8, and 68.3 ± 1.2 °C, respectively.

### Chemicals and reagents

All chemicals were purchased from Sigma-Aldrich (MO, USA) and/or Thermo Fisher Scientific (MA, USA), unless specified elsewhere. For molecular cloning, restriction enzymes and T4 ligase were obtained from New England Biolabs (MA, USA). Phusion Hot Start II DNA polymerase was used for polymerase chain reaction (PCR).

### Media and cultivation

For molecular cloning and protein expression, *E. coli* strains were grown in lysogeny broth (LB) containing appropriate antibiotics unless noted otherwise. For *in vivo* characterization of CAT_Sa_ in *E. coli*, M9 hybrid medium [5] with 20 g/L glucose was used. For *C. thermocellum* culture, MTC minimal medium or CTFuD-NY medium [30] was used as specified in the experiments. Optical density (OD) was measured by a spectrophotometer at 600 nm wavelength (Spectronic 200+, Thermo Fisher Scientific, MA, USA).

### Multiple sequence alignment analysis

Multiple sequence alignment (MSA) analysis was performed using MEGA7 [31]. Protein sequences were aligned by ClustalW [32] and visualized by ESPript 3.0 (http://espript.ibcp.fr) [33]. The key features in protein structures of 3U9F [34], 4CLA [35], and 2XAT [36] were extracted from CAT_SALTI, CAT3_ECOLIX, and CAT4_PSEAE, respectively.

### Molecular modeling and docking simulations

#### Three-dimensional (3D) structures

The 3D structure of CAT_Sa_ and alcohols of interest were first generated using Swiss-Model [37] and the ‘Builder’ tools of MOE (Molecular Operating Environment software, version 2019.01), respectively. The 3D structure of the dual substrates-bounded CAT_Sa_ complex (i.e., acetyl-CoA-isobutanol-CAT_Sa_) was obtained by extracting an isobutanol from the isobutanol-CAT_Sa_ complex and then adding it to the acetyl-CoA-CATSa complex. All the structures were prepared by the ‘QuickPrep’ tool of MOE with default parameters and further optimized by energy minimization with the Amber10: EHT force field.

#### Docking simulation

To perform docking simulations, the potential binding pocket was searched using the ‘Site Finder’ tool of MOE. The best-scored site, consistent with the reported catalytic sites [38], was selected for further studies. Docking simulations were performed as previously described [39]. Briefly, acetyl-CoA and each alcohol were docked using the induced fit protocol with the Triangle Matcher placement method and the London ΔG scoring function. After the docking simulations, the best-scored binding pose, showing the crucial interaction between the residue and the substrate at root-mean-square-deviation (RMSD) < 2 Å, was selected. As an example, for the acetyl-CoA docking, the binding pose exhibiting the hydrogen bond between the hydroxyl of Ser-148 and the N^71^ of the CoA was chosen [40]. For the alcohol docking, the binding pose showing the hydrogen bond between the N^3^ of His-189 and the hydroxyl of alcohol was selected [22].

#### In silico mutagenesis analysis

*In silico* mutagenesis analysis of the acetyl-CoA-isobutanol-CAT_Sa_ complex was carried out as previously described [39]. Specifically, the ‘alanine scan’ and ‘residue scan’ tools of MOE were used to identify the potential residue candidates for mutagenesis.

### Molecular cloning

#### Plasmid construction

Plasmids were constructed by the standard molecular cloning technique of ligase dependent method and/or Gibson assembly [41] using the primers listed in Table S1. The constructed plasmids were introduced into *E. coli* TOP10 by heat shock transformation. Colonies isolated on a selective plate were PCR screened and plasmid purified. The purified plasmids were verified via Sanger sequencing before being transformed into *E. coli* BL21 (DE3). Site-directed mutagenesis was performed using the QuickChange™ site-directed mutagenesis protocol with reduced overlap length [42] or Gibson assembly method [41]. For the *C. thermocellum* engineering, the plasmid pHS005 was constructed first and then modified to pHS0024. pHS0024 has no *hpt* at the downstream of the operon while other sequences of the plasmid are identical to pHS005.

#### Transformation

The conventional chemical transformation and electroporation methods were used for transformation of *E. coli* [43] and *C. thermocellum* [30], respectively. For *C. thermocellum*, the method, however, was slightly modified as described here. First, *C. thermocellum* M1354 (Table 1) was cultured in 50 mL CTFuD-NY medium at 50°C inside an anaerobic chamber (Bactron300, Sheldon manufacturing Inc., OR, USA). The cell culture with OD in a range of 0.8-1.0 was cooled down at room temperature for 20 minutes. Beyond this point, all steps were performed outside the chamber. The cooled cells were harvested at 6,500 x g and 4°C for 20 minutes. The cell pellets were washed twice with ice-chilled Milli-Q water and resuspended in 200 μL of the transformation buffer consisting of 250 mM sucrose and 10% (v/v) glycerol. Several 30 μL aliquots of the electrocompetent cells were immediately stored at −80°C for further use. For electroporation, the electrocompetent cells were thawed on ice and incubated with 500−1,000 ng of methylated plasmids [44] for 10 minutes. Then, the cells were transferred to an ice-chilled 1-mm gap electroporation cuvette (BTX Harvard Apparatus, MA, USA) followed by two consecutive exponential decay pulses with 1.8 kV, 350 Ω, and 25 μF. The pulses usually resulted in a 7.0-8.0 ms time constant. The cells were immediately resuspended in pre-warmed fresh CTFuD-NY and recovered at 50°C under anaerobic condition (90 % N_2_, 5% H_2_, and 5% CO_2_) inside a rubber capped Balch tube. After 0-12 hours of recovery, the cells were mixed with molten CTFuD-NY agar medium supplemented with 15 μg/mL thiamphenicol. Finally, the medium-cell mixture was poured on a petri dish and solidified inside the anaerobic chamber. The plate was incubated at 50°C up to one week until colonies appeared.

### *In vivo* characterization of CAT_Sa_ and its variants in *E. coli*

For *in vivo* characterization of CAT_Sa_ and its variants in *E. coli*, high-cell density cultures were performed as previously described [45] with an addition of 2 g/L of various alcohols. For *in-situ* extraction of esters, each tube was overlaid with 25% (v/v) hexadecane. To confirm the protein expression of CAT_Sa_ and its variants, 1% (v/v) of stock cells were grown overnight at 37°C and 200 rpm in 15 mL culture tubes containing 5 mL of LB media and antibiotics. Then, 4% (v/v) of the overnight cultures were transferred into 1 mL of LB media containing antibiotics in a 24-well microplate. The cultures were grown at 37°C and 350 rpm using an incubating microplate shaker (Fisher Scientific, PA, USA) until OD reached to 0.4∼0.6 and then induced by 0.1 mM isopropyl β-D-1-thiogalactopyranoside (IPTG) for 4 hours with a Breathe-Easy Sealing Membrane to prevent evaporation and cross contamination (cat# 50-550-304, Research Products International Corp., IL, USA). The protein samples were obtained using the B-PER complete reagent (cat# 89822, Thermo Scientific, MA, USA), according to the manufacturer’s instruction and analyzed by SDS-PAGE.

### Enzyme characterization

#### His-tag purification

For enzyme expression, an overnight culture was inoculated with a 1:50 ratio in fresh LB medium containing 1 mM IPTG and antibiotics, followed by 18°C overnight incubation (up to 20 hours) in a shaking incubator at 200 rpm. The induced cells were harvested by centrifugation at 4°C, and 4,700 x g for 10 minutes. The cell pellet was then washed once with Millipore water and resuspended in the B-PER complete reagent. After 30 min incubation at room temperature, the mixture was centrifuged at 17,000 x g for 2 minutes. The supernatant was collected and designated as crude extract. For his-tag purification, the crude extract was incubated with HisPur Ni-NTA superflow agarose in a batch as the manufacturer recommends. Then, the resin was washed with at least three volumes of wash buffer, consisting of 50 mM Tris-HCl (pH 8.0), 300 mM NaCl, 10 mM imidazole, and 0.1 mM EDTA. The resin bound proteins were eluted by 300 μL elution buffer containing 50 mM Tris-HCl (pH 8.0), 50 mM NaCl, 300 mM imidazole, and 0.1 mM EDTA. The eluted sample was then desalted and concentrated via an Amicon filter column with 10 kDa molecular weight cutoff. Finally, the protein sample was suspended in 200 μL of 20 mM Tris-HCl buffer (pH 8.0). Protein concentration was measured by the Bradford assay [46] with bovine serum albumin (BSA) as the reference protein.

#### Thermal shift assay

To measure protein melting point (Tm), a thermofluor assay was employed with SYPRO Orange [47]. About 10 to 250 μg of His-tag purified protein was mixed with 5x SYPRO Orange in a 50 μL final volume in a 96-well qPCR plate. The plate was sealed with PCR caps before running the assay. The StepOne real-time PCR machine (Applied Biosystems, CA, USA) was used to run the assay with the following parameters: ROX reporter, 1°C increment per cycle, one-minute hold at every cycle, and temperature range from 20°C to 98°C. The data was collected, exported, and processed to calculate Tm.

#### 5,5’-dithiobis-(2-nitrobenzoic acid) (DTNB) assay

Reaction rate for each CAT was determined by a DTNB assay [48] in a 384-well plate. Total reaction volume was 50 μL with the reaction buffer comprising of 50 mM Tris-HCl (pH 8.0). Concentrations of acetyl-CoA (CoALA Biosciences, TX, USA) and alcohols were varied as specified in each experiment. Final enzyme concentrations of 0.05 μg/mL and 10 μg/mL were used for the reactions towards chloramphenicol and alcohols, respectively. Reaction kinetics were collected by measuring absorbance at 412 nm every minute for one hour at 50°C in a microplate reader (Synergy HTX microplate reader, BioTek). The reaction rate was calculated using the extinction coefficient from a standard curve of free coenzyme A (MP Biomedicals, OH, USA) under the same condition. It should be noted that since the maximum operating temperature recommended for the plate reader is 50°C, the high throughput enzyme assay for CAT at elevated temperature was only performed to determine enzyme kinetics parameters.

#### Calculation of kinetic parameters for reaction rates

The parameters of Michaelis-Menten rate law (eqn. 1) were calculated for each enzyme as follows. First, linear regression was performed on data collected from a microplate reader to identify initial reaction rates, *y_i_*, at different initial substrate concentrations, *s_i_*, where *i= {1,2,…,n}* is the number of data points collected. Then, these initial reaction rates and associated initial substrate concentrations for all replicates were simultaneously fit to the Michaelis-Menten model (eqn. 1) using robust non-linear regression (eqn. 2) with a soft-L1-loss estimator (eqn. 3) as implemented in the SciPy numerical computing library v1.2.0 [49, 50].

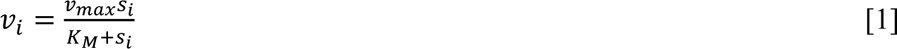

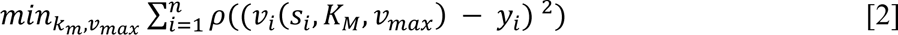

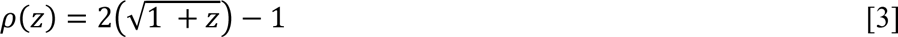

The least squares problem determines the parameters *K_M_* and *υ_max_* by minimizing the difference between the model predicted reaction rates *υ_i_* and measured reaction rates *y_i_* (eqn. 2). A smoothing function *ρ*(*z*) is used to make the least square problem resistant to outliers (eqn. 3). Due to the unbiased resistance to outliers and the avoidance of errors resulting from conventional linearization methods, robust non-linear regression provides the most precise parameter estimate for the Michaelis-Menten model [51].

### Isobutyl acetate production in *C. thermocellum*

#### Cellobiose fermentation

Isobutyl acetate production from cellobiose in *C. thermocellum* strains was performed by the two-step bioconversion configuration. Cells were first cultured in MTC minimal medium [30] containing 5 g/L cellobiose in a rubber capped Balch tube until OD reached 0.8∼1.0. The cells were cooled down at room temperature for 20 minutes and centrifuged at 4,700 x g and 4°C for 20 minutes. After removing the supernatant, cells were resuspended in the same volume of fresh MTC minimal media containing 2 g/L isobutanol in an anaerobic chamber. The cell suspension was then divided into 800 μL in a 2.0 mL screw cap microcentrifuge tube with a 200 μL hexadecane overlay. The cells were incubated at 55°C for 24 hours followed by analysis of gas chromatography coupled with a mass spectrometer (GC/MS) to quantify the amount of isobutyl acetate produced.

#### Cellulose fermentation

For the cellulose fermentation, modified MTC medium (C-MTC medium) was used. 20 g/L of Avicel PH-101 was used as a sole carbon source instead of cellobiose, and 10 g/L of MOPS was added to increase buffer capacity. Initial pH was adjusted to 7.5 by 5M KOH and autoclaved. In an anaerobic chamber, 0.8 mL of overnight cell culture was inoculated in 15.2 mL of C-MTC medium (1:20 inoculation ratio) with 4 mL of overlaid hexadecane. Each tube contained a small magnetic stirrer bar to homogenize cellulose. The rubber capped Balch tube was incubated in a water bath connected with a temperature controller set at 55°C and a magnetic stirring system. Following pH adjustment with 70 μL of 5 M KOH injection, 800 μL of cell culture and 200 μL of hexadecane layer were sampled every 12 hours. Culture pH was maintained within a range of 6.4-7.8 during the fermentation.

Cell growth was monitored by measuring pellet protein. The cell-cellulose pellet from 800 μL sampling volumes was washed twice with Milli-Q water and suspended by 200 μL lysis buffer (0.2 M NaOH, 1% SDS) followed by an hour incubation at room temperature. Then, the solution was neutralized with 50 μL 0.8 M HCl and diluted by 550 μL water. The mixture was centrifuged at 17,000 x g for 3 minutes. Protein concentration from the supernatant was analyzed by the detergent-compatible Bradford assay (Thermo Scientific, WA, USA). The residual pellet was boiled in a 98°C oven for an hour before quantifying residual cellulose.

Residual cellulose was quantified by the phenol-sulfuric acid method [52] with some modifications. The boiled sample was washed twice with Milli-Q water and suspended in 800 μL water to make equivalent volume to the original. The sample was homogenized by pipetting and vortexing for 10 seconds, and 20 μL of the homogenized sample was transferred to a new 2.0 mL microcentrifuge tube or 96-well plate and dried overnight in a 55°C oven. The dried pellet was suspended in 200 uL of 95% sulfuric acid and incubated for an hour at room temperature. After the pellet was dissolved completely, 20 μL of 5% phenol was added and mixed with the sulfuric acid solution. After 30 min incubation at room temperature, 100 μL of the sample was transferred to a new 96-well plate, and the absorbance at 490 nm was measured. The absorbance was converted to cellulose concentration by the standard curve of Avicel PH-101 treated by the same procedure.

### Analytical methods

#### High-performance liquid chromatography (HPLC)

Extracellular metabolites were quantified by using a high-performance liquid chromatography (HPLC) system (Shimadzu Inc., MD, USA). 800 μL of culture samples was centrifuged at 17,000 x g for 3 minutes, then the supernatants were filtered through 0.2 micron filters and run with 10 mN H_2_SO_4_ mobile phase at 0.6 mL/min on an Aminex HPX-87H (Biorad Inc., CA, USA) column at 50°C. Refractive index detector (RID) and ultra-violet detector (UVD) at 220 nm were used to monitor concentrations of sugars, organic acids, and alcohols.

#### Gas chromatography coupled with mass spectroscopy (GC/MS)

Esters were measured by GC (HP 6890, Agilent, CA, USA) equipped with a MS (HP 5973, Agilent, CA, USA). For the GC system, the Zebron ZB-5 (Phenomenex, CA, USA) capillary column (30 m x 0.25 mm x 0.25 μm) was used to separate analytes, and helium was used as the carrier with a flow rate of 0.5 mL/min. The oven temperature program was set as follows: 50°C initial temperature, 1°C/min ramp up to 58°C, 25°C/min ramp up to 235°C, 50°C/min ramp up to 300°C, and 2-minutes bake-out at 300°C. 1 μL of sampled hexadecane layer was injected into the column in the splitless mode with an injector temperature of 280°C. For the MS system, selected ion mode (SIM) was used to detect and quantify esters with the following parameters: (i) ethyl acetate, m/z 45.00 and 61.00 from 4.2 to 4.6 minute retention time (RT), (ii) isopropyl acetate, m/z 45 and 102 from 4.7 to 5.0 minute RT, (iii) propyl acetate, m/z 59 and 73 from 5.2 to 5.8 minute RT, (iv) ethyl isobutyrate, m/z 73 and 116 from 6.1 to 6.6 minute RT, (v) isobutyl acetate, m/z 61 and 101 from 6.6 to 7.6 minute RT, (vi) butyl acetate, m/z 61 and 116 from 7.7 to 9.2 minute RT, (vii) isobutyl isobutyrate, m/z 89 and 129 from 10.1 to 12.5 minute RT, (viii) benzyl acetate, m/z 108 and 150 from 13.1 to 13.8 minute RT, and (ix) 2-phenethyl acetate, m/z 104 and 121 from 13.8 to 15.5 minute RT. Isoamyl alcohol and isoamyl acetate were used as the internal standard analytes. The esters were identified by RT and quantified by the peak areas and standard curves. Standard curves were determined by using pure esters diluted into hexadecane at concentrations of 0.01 g/L, 0.05 g/L, 0.1 g/L, 0.5 g/L, and 1 g/L.

## ABBREVIATIONS

AAT: alcohol acetyltransferase
CBP: consolidated bioprocessing
CAT: chloramphenicol acetyltransferase
PCR: polymerase chain reactions
MSA: multiple sequence alignment
DCW: dried cell weight
DTNB: 5,5’-dithiobis-(2-nitrobenzoic acid)
GC: gas chromatography
HPLC: high-performance liquid chromatography
IPTG: isopropyl β-D-1-thiogalactopyranoside
kDa: kilo Dalton
MOE: Molecular Operating Environment software
MS: mass spectrometry
OD: optical density
RMSD: root-mean-square-deviation
RT: retention time
SDS-PAGE: sodium dodecylsulfate polyacrylamide gel electrophoresis
8-AZH: 8-Azahypoxanthine
Tm: melting point

## AUTHOR’S CONTRIBUTIONS

CTT initiated and supervised the project. HS, JWL and CTT designed the experiments, analyzed the data, and drafted the manuscript. HS and JWL performed the experiments. SG calculated enzyme kinetic parameters and edited the manuscript. All authors read and approved the final manuscript.

## ACKNOWLEDGMENTS

The authors would like to thank the Center of Environmental Biotechnology at UTK for using the GC/MS instrument. We would also like to acknowledge the gene synthesis from The Joint Genome Institute.

## COMPETING INTERESTS

The authors declare that they have no competing interests.

## AVAILABILITY OF SUPPORTING DATA

One additional file contains supporting data.

## CONSENT FOR PUBLICATION

All the authors consent for publication.

## ETHICAL APPROVAL AND CONSENT TO PARTICIPATE

Not applicable.

## FUNDING

This research was financially supported in part by the NSF CAREER award (NSF#1553250) and the Center for Bioenergy Innovation (CBI), the U.S. Department of Energy (DOE) Bioenergy Research Centers funded by the Office of Biological and Environmental Research in the DOE Office of Science. The work conducted by the U.S. Department of Energy Joint Genome Institute, a DOE Office of Science User Facility, is supported by the Office of Science of the U.S. Department of Energy under Contract No. DE-AC02-05CH11231.

