## Supplementary Materials for "Single mutation at a highly conserved region of chloramphenicol acetyltransferase enables thermophilic isobutyl acetate production directly from cellulose by *Clostridium thermocellum*"

Running title: Engineering of a thermostable chloramphenicol acetyltransferase for isobutyl acetate production at elevated temperature

**Table S1.** List of primers used in this study. The bold and underlined letters indicate restriction and site-directed mutation sites, respectively.

| Primers | Primer sequence (5' to 3') |
|---|---|
| ***Site-saturation mutagenesis of CAT$_{Sa}$ F97*** | |
| CAT$_{Sa}$_F BamHI | CTCT**GGATCC**AATGAACTTTAATAAAATTGATTTAG |
| CAT$_{Sa}$_R SacI | CTCT**GAGCTC**TTATAAAAGCCAGTCATTAGGCCTA |
| F97R_F | TGATGGTGTATCTAAAACACGTTCTGGTATTTGGACTC |
| F97R_R | GTCCAAATACCAGAACGTGTTTTAGATACAC |
| F97K_F | TGATGGTGTATCTAAAACAAAATCTGGTATTTGGACTC |
| F97K_R | GTCCAAATACCAGATTTTGTTTTAGATACAC |
| F97L_F | TGATGGTGTATCTAAAACACTGTCTGGTATTTGGACTC |
| F97L_R | GTCCAAATACCAGACAGTGTTTTAGATACAC |
| F97W_F | TGATGGTGTATCTAAAACATGGTCTGGTATTTGGACTC |
| F97W_R | GTCCAAATACCAGACCATGTTTTAGATACAC |
| F97I_F | TGATGGTGTATCTAAAACAATTTCTGGTATTTGGACTC |
| F97I_R | GTCCAAATACCAGAAATTGTTTTAGATACAC |
| F97G_F | TGATGGTGTATCTAAAACAGGTTCTGGTATTTGGACTC |
| F97G_R | GTCCAAATACCAGAACCTGTTTTAGATACAC |
| CAT$_{Sa}$_F | TGTTTAACTTTAAGAAGGAGATATACCAT |
| BB_CAT$_{Sa}$_R | TGGATCCTGGCTGTGG |
| F97A_R | CATTCTTTACAGGAGTCCAAATACCAGAGGCTGTTTTAGATACACCATCAAAAATTGTAT |
| BB_F97A_F | GCCTCTGGTATTTGGACTCCTGTA |
| F97N_R | CATTCTTTACAGGAGTCCAAATACCAGAATTTGTTTTAGATACACCATCAAAAATTGTAT |
| BB_F97N_F | AATTCTGGTATTTGGACTCCTGTA |
| F97D_R | CATTCTTTACAGGAGTCCAAATACCAGAGTCTGTTTTAGATACACCATCAAAAATTGTAT |
| BB_F97D_F | GACTCTGGTATTTGGACTCCTGTA |
| F97C_R | CATTCTTTACAGGAGTCCAAATACCAGAACATGTTTTAGATACACCATCAAAAATTGTAT |
| BB_F97C_F | TGTTCTGGTATTTGGACTCCTGTA |
| F97Q_R | CATTCTTTACAGGAGTCCAAATACCAGATTGTGTTTTAGATACACCATCAAAAATTGTAT |
| BB_F97Q_F | CAATCTGGTATTTGGACTCCTGTA |
| F97E_R | CATTCTTTACAGGAGTCCAAATACCAGACTCTGTTTTAGATACACCATCAAAAATTGTAT |
| BB_F97E_F | GAGTCTGGTATTTGGACTCCTGTA |
| F97H_R | CATTCTTTACAGGAGTCCAAATACCAGAGTGTGTTTTAGATACACCATCAAAAATTGTAT |
| BB_F97H_F | CACTCTGGTATTTGGACTCCTGTA |
| F97M_R | CATTCTTTACAGGAGTCCAAATACCAGACATTGTTTTAGATACACCATCAAAAATTGTAT |
| BB_F97M_F | ATGTCTGGTATTTGGACTCCTGTA |
| F97P_R | CATTCTTTACAGGAGTCCAAATACCAGATGGTGTTTTAGATACACCATCAAAAATTGTAT |
| BB_F97P_F | CCATCTGGTATTTGGACTCCTGTA |
| F97S_R | CATTCTTTACAGGAGTCCAAATACCAGAGCTTGTTTTAGATACACCATCAAAAATTGTAT |
| BB_F97S_F | AGCTCTGGTATTTGGACTCCTGTA |
| F97T_R | CATTCTTTACAGGAGTCCAAATACCAGACGTTGTTTTAGATACACCATCAAAAATTGTAT |
| BB_F97T_F | ACGTCTGGTATTTGGACTCCTGTA |
| F97Y_R | CATTCTTTACAGGAGTCCAAATACCAGAGACTGTTTTAGATACACCATCAAAAATTGTAT |
| BB_F97Y_F | GTCTCTGGTATTTGGACTCCTGTA |
| F97V_R | CATTCTTTACAGGAGTCCAAATACCAGAGTATGTTTTAGATACACCATCAAAAATTGTAT |
| BB_F97V_F | TACTCTGGTATTTGGACTCCTGTA |
| ***Clostridium thermocellum engineering*** | |
| *pHS005 construction (C. thermocellum markless gene deletion plasmid)* | |
| pNW33N backbone F | GAGGGGTTTTTTGC CAATCCCGTTTGTTGAACTAC |
| pNW33N backbone R | CAGGAAACAGCTATGA CAGGAAACAGCTATGACCATGA |
| PgapDH F | TCATAGCTGTTTCCTGTAATTACTGTATCTCTCTGGC |

| | |
|---|---|
| PgapDH R | AATTTTATTAAAGTTCATTAATATCGCCTCCTATTGTAA |
| $CAT_{Sa}$ F | AGGAGGCGATATTAATGAACTTTAATAAAATTGATTTAGACAATTGG |
| $CAT_{Sa}$ R | ATCATGACGTCGACCTCCTTTATTATAAAAGCCAGTCATTAGGCCTATC |
| hpt F | AAAGGAGGTCGACGTCATGATAAATCAAATTAAAGAAATTTTGG |
| hpt R | CGGCCGTGTACAATAGCAAAACACTATCTCTCATAC |
| 2927 terminator F | GACAAAGAAGATATGGACTAAAAAATATACAAAGGTTTCTTG |
| 2927 terminator R | GATTATGCGGCCGTGTACAATAGCAAAACACTATCTCTCATACA |
| MCS2 F | ATTGTACACGGCCGCATAATC |
| MCS2 R | CAAACGGGATTGGCAAAAAACCCCTCAAGACC |
| bb tdk F | TTTCCCGTTCTCTCTGATTGTGA |
| bb tdk R | GCAACTATGGATGAACGAAATAGA |
| tdk F | ATTTCGTTCATCCATAGTTGCAAAAATCCGCTTAAGTCCGCG |
| tdk R | CAATCAGAGAGAACGGGAAAGTTTCCGTATAAATTAACCGTATG |
| *pHS0024 construction (pHS005 without hpt gene)* | |
| -hpt F | AAAATATACAAAGGTTTCTTGTG TTTTTAATACCGTTATGTTAATATAATG |
| -hpt R | CACAAGAAACCTTTGTATATTTT TTATAAAAGCCAGTCATTAGGC |

**Table S2.** $K_M$ values of $CAT_{Sa}$ and $CAT_{Sa}$ F97W towards acetyl-CoA.

| | $CAT_{Sa}$ | | $CAT_{Sa}$ F97W | |
|---|---|---|---|---|
| Co-substrates | Chloramphenicol | Isobutanol | Chloramphenicol | Isobutanol |
| $K_M$ (mM) | 0.08 ± 0.01 | 0.06 ± 0.01 | 0.09 ± 0.01 | 0.08 ± 0.02 |

**Figure S1. (A)** Reaction mechanisms of acetylation of chloramphenicol by $CAT_{Sa}$. Briefly, the acetylation of chloramphenicol involves three steps. First, the 3-hydroxyl group of the chloramphenicol is deprotonated by the imidazole ring of histidine of the CAT's active site, generating the activated oxygen. Then, the nucleophilic attack by the oxyanion at the thioester carbonyl carbon of acetyl-CoA generates a tetrahedral intermediate. Finally, loss of the free CoA yields the 3-acetylchloramphenicol. **(B)** Physical properties of various alcohols used in this study and the predicted $\Delta G_{bind}$ from the docking simulation. **(C)** Correlation between the predicted $\Delta G_{bind}$ and the molecular weights of alcohols. **(D)** Correlation between the predicted $\Delta G_{bind}$ and the LogP values of alcohols.

A

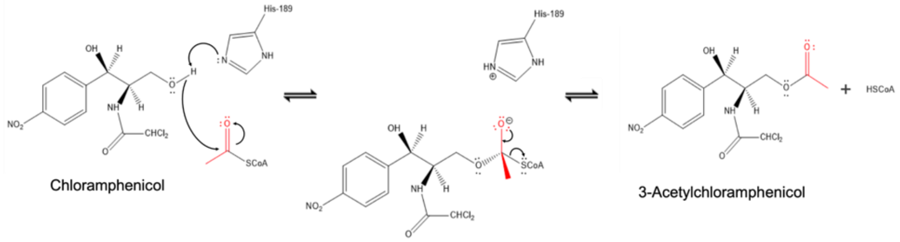


B

| Compound | Molecular formula | Structural formula | Molecular weight (g/mol)* | LogP* | Predicted $\Delta G_{bind}$ (kcal/mol) |
|---|---|---|---|---|---|
| Ethanol | $C_2H_6O$ | 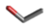 | 46.069 | -0.31 | -3.28 |
| Propanol | $C_3H_8O$ | 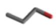 | 60.096 | 0.34 | -3.56 |
| Isopropanol | $C_3H_8O$ | 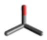 | 60.096 | 0.05 | -3.17 |
| Butanol | $C_4H_{10}O$ | 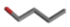 | 74.123 | 0.88 | -3.78 |
| Isobutanol | $C_4H_{10}O$ | 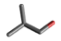 | 74.123 | 0.76 | -3.84 |
| Benzyl alcohol | $C_7H_8O$ | 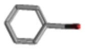 | 108.14 | 1.10 | -4.03 |
| 2-Phenethyl alcohol | $C_8H_{10}O$ | 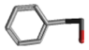 | 122.16 | 1.36 | -4.16 |
| Chloramphenicol | $C_{11}H_{12}C_{l2}N_2O_5$ | 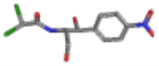 | 323.126 | 1.14 | -6.49 |

*Reference: Hansch, C., Leo, A., Hoekman, D. H., 1995. Exploring QSAR. American Chemical Society, Washington, DC.

C

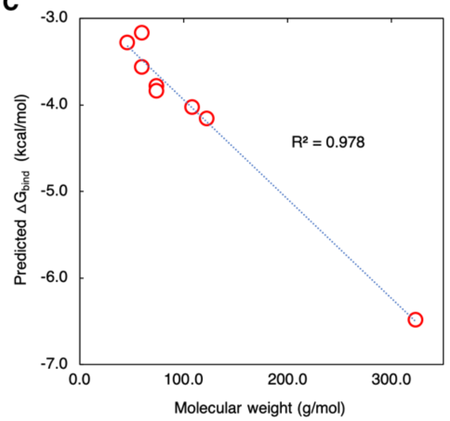


D

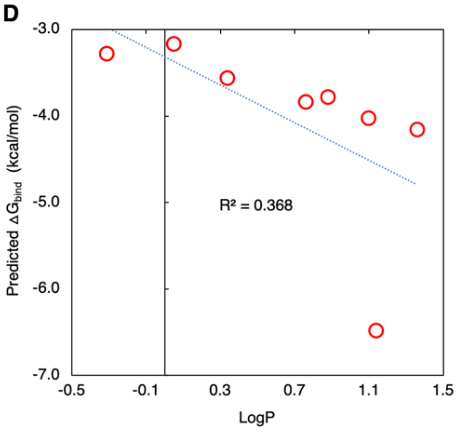

**Figure S2.** Multiple sequence alignment of CATs. **(A)** Sequence alignment of 22 CATs classified as Type A. **(B)** Sequence alignment of 27 CATs including Type A and Type B. The highly conserved regions are red highlighted.

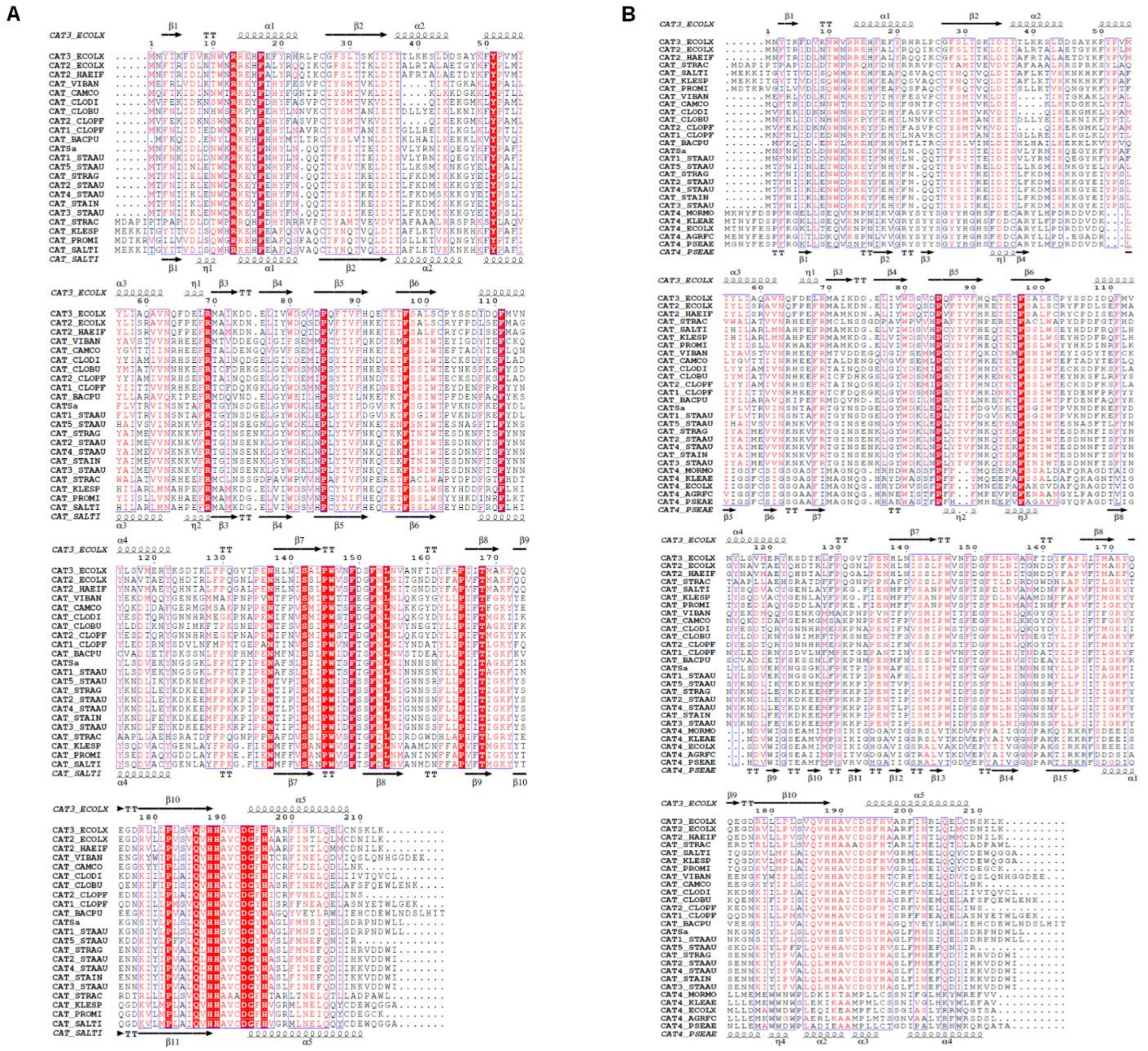

**Figure S3. (A)** Alanine scan of 23 identified residues in the binding pocket of the $CAT_{Sa}$-isobutanol-acetyl-CoA complex. ΔStability (kcal/mol) represents the relative stability of each variant with respect to its wild type. Positive ΔStability values indicate that the structural stability of the protein-ligands complex is reduced if a specific residue is replaced with alanine, suggesting the significance of that specific residue in the protein-ligands complex. **(B)** Representative residue scan from a selection of six selected residues identified from alanine scan in the binding pocket of the $CAT_{Sa}$-isobutanol-acetyl-CoA or $CAT_{Sa}$-chloramphenicol-acetyl-CoA complex. To identify the significant residues in the isobutanol binding exclusively, the $\Delta\Delta Stability_{(Cm\text{-}iBtOH)}$ was calculated by subtracting $\Delta Stability_{iBtOH}$ from $\Delta Stability_{Cm}$.

**A**

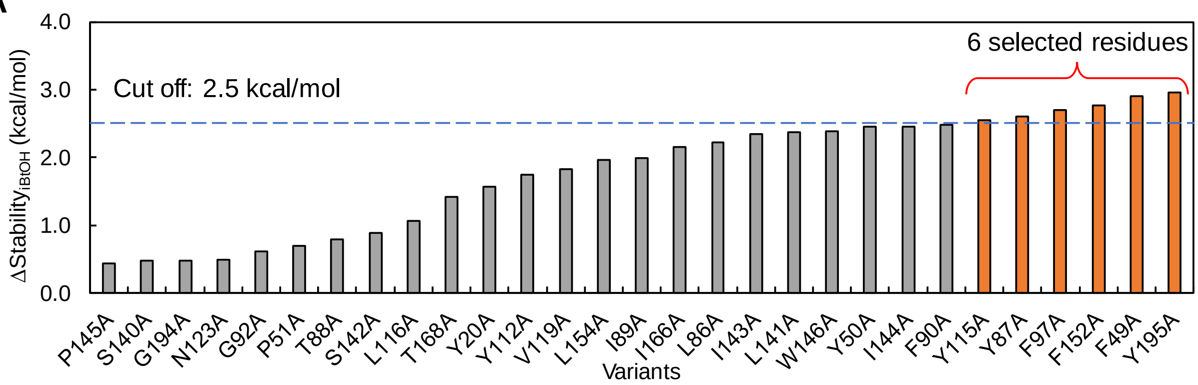


**B**

| Rank | Variants | $\Delta Stability_{Cm}$ | $\Delta Stability_{iBtOH}$ | $\Delta\Delta Stability_{(Cm\text{-}iBtOH)}$ |
|---|---|---|---|---|
| 1 | F97Y | 2.32 | 1.02 | 1.30 |
| 2 | F97W | 2.15 | 0.89 | 1.26 |
| 3 | F97V | 2.70 | 1.67 | 1.03 |
| 4 | Y195E | 2.79 | 1.94 | 0.85 |
| 5 | Y195N | 2.69 | 1.85 | 0.83 |
| 6 | F97H | 2.63 | 1.98 | 0.65 |
| 7 | Y87G | 4.51 | 3.90 | 0.61 |
| 8 | F152W | 0.90 | 0.37 | 0.53 |
| 9 | Y195R | 2.28 | 1.81 | 0.47 |
| 10 | Y195Q | 2.13 | 1.68 | 0.45 |
| 11 | Y115K | 1.86 | 1.51 | 0.35 |
| 12 | F97C | 2.69 | 2.43 | 0.27 |
| 13 | F152E | 2.51 | 2.25 | 0.26 |
| 14 | Y195M | 2.14 | 1.89 | 0.25 |
| 15 | F152H | 2.21 | 1.96 | 0.25 |

Cut off: 1.0 kcal/mol (between ranks 3 and 4)

**Figure S4.** An overlaid GC/MS chromatogram showing the trace amount of isobutyl acetate produced by the wildtype *C. thermocellum.*

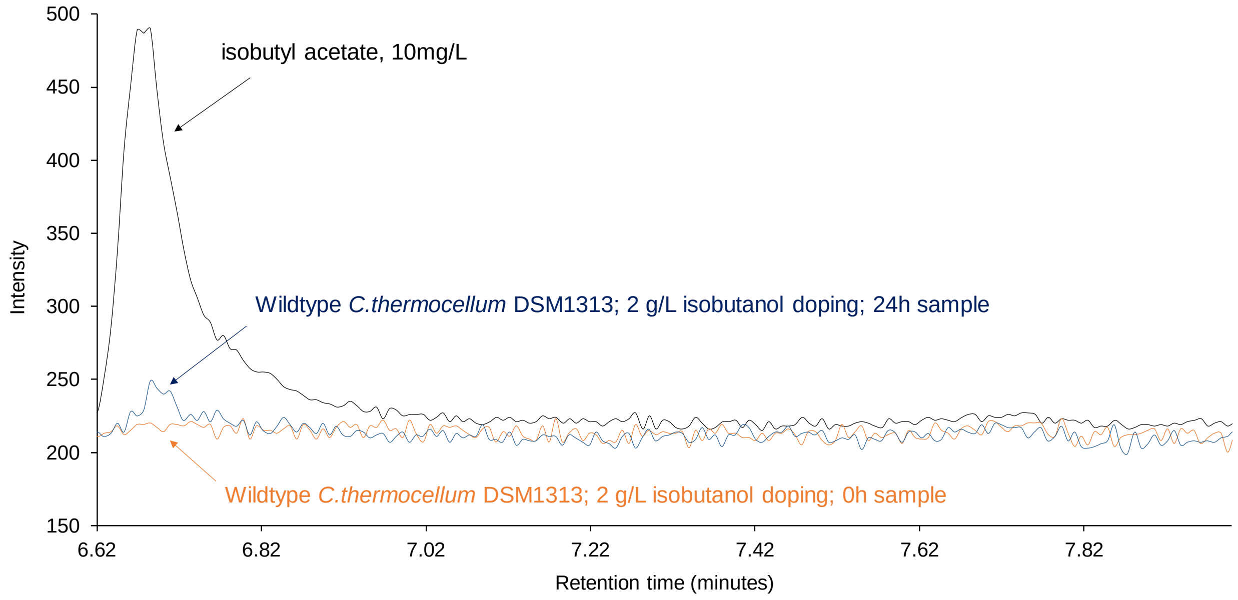

**Figure S5.** Fementative products from cellulose fermentation of HSCT0102. **(A)** Simplified metabolic pathways of C. thermocellum for biosynthesis of native fermentative metabolites and heterologous esters. It should be noted that *C. thermocellum* has two unique routes to endogenously produce isobutanol from 2-ketoisovalerate. The first route, that was experimentally validated [1], uses the 2-ketoisovalerate ferredoxin oxidoreductase (KOR) and alcohol dehydrogenase (ADH) converts 2-ketoisovalerate into isobutyryl-CoA and then isobutanol. The second route, that has not yet been experimentally validated, uses keto-isovalerate decarboxylase (KIVD) and ADH to covert 2-ketoisovalerate to isobutyraldehyde and then isobutanol. Heterologous, thermostable KIVD was engineered in *C. thermocellum* [1]; however, the endogenous acetolactate synthase (*ilvBN*), that catalyzes the first step in valine biosynthesis and might have KIVD activity as observed in *E. coli* [2, 3], has not yet been experimentally confirmed in *C. thermocellum*. **(B)** Kinetic profile of biosynthesis of other esters besides isobutyl acetate, including ethyl acetate, ethyl isobutyrate, and isobutyl butyrate. **(C)** An overlaid GC/MS chromatogram demonstrating ester biosynthesis. **(D)** Kinetic profile of other fermentative metabolites. Abbreviation: IS, internal standard.

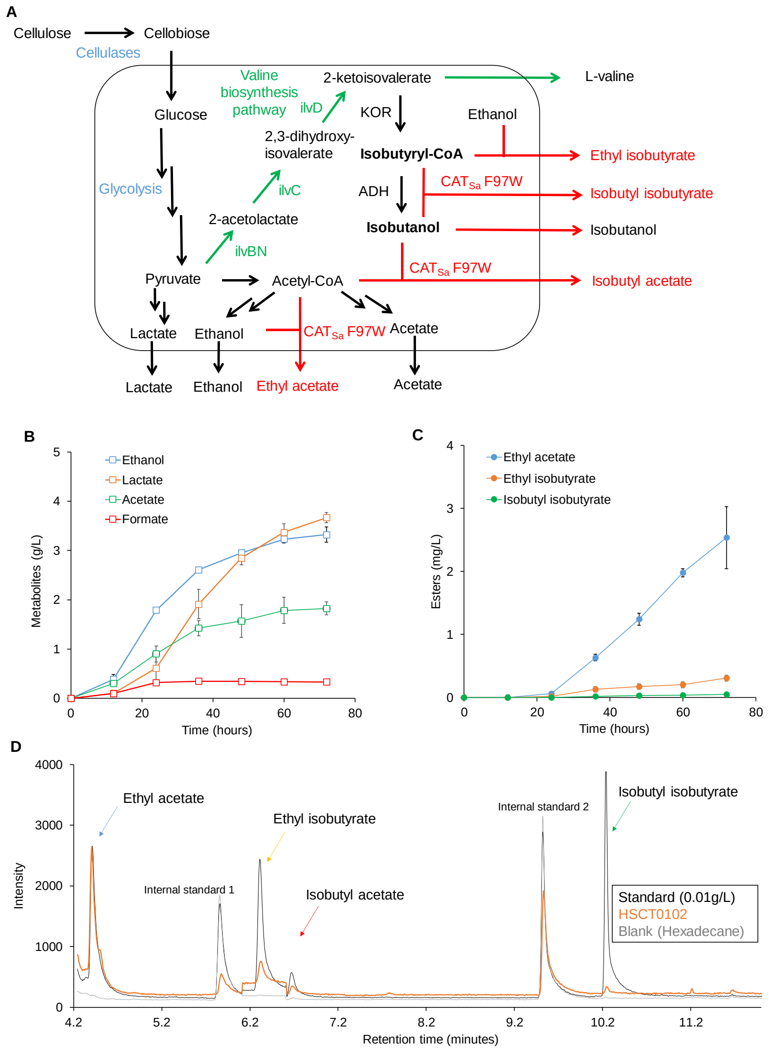
